# Age, not body size, influences flight distance in the bumble bee *Bombus terrestris* L. (Hymenoptera: Apidae)

**DOI:** 10.1101/2024.02.07.579366

**Authors:** Milena Gilgenreiner, Christoph Kurze

## Abstract

Bumble bees and honey bees provide crucial pollination service and have become important insect model system. Despite their close relation, they differ in their morphology, physiology, and social organisation. Notably, honey bees workers exhibit age-based polyethism, while bumble bees display considerable body size variation. Though body size is known to affect foraging range, behaviour, and flight ability, the influence of age remains less explored. Here we studied the flight performance (distance and speed) in aging bumble bee workers using tethered flight mills. Additionally, we measured their intertegular distance (ITD), dry mass, and fat content. Flight distance was predominantly influenced by age, challenging assumptions that age does not play a role in foraging and task allocation. Between 7 to 14 days, flight distance increased six-fold before a slight decline at the age of 21 days, aligning with age-dependent flight physiology in honey bees. Larger workers had decreasing mass-specific fat reserves, challenging the notion about their energy preservation for oocyte development. Our findings indicate that age substantially influences flight distance, impacting foraging performance and potentially altering task allocation strategies. This underscores the need to consider individual age and physiological changes alongside body size and mass in experiments involving bumble bee workers.

## 1. Introduction

Division of labour, or better task allocation, is fundamental to the ecological success of eusocial insects such as ants and social bees [1]. Eusociality, by definition, requires a division between reproductive and non-reproductive tasks, with a minority engaging in reproduction and the majority comprising workers with little or no direct reproduction. Workers primarily perform tasks such as brood care, nest construction and maintenance as well as foraging [2, 3]. Age-dependent behaviour (age-based polyethism) or morphological variation are often considered the underlying mechanism of task allocation [4].

Age-based polyethism has been well studied in the honey bees *Apis mellifera* (Hymenoptera: Apidae). Young workers perform in-hive tasks, including cleaning, brood care, comb building, and food processing for the first weeks of their adult life. As workers age, they transition to out-hive tasks, including guarding duties and nectar or pollen foraging. These behavioural transitions are mediated by physiological changes, including alterations in juvenile hormone (JH) [5]. Hormonal levels can also be modulated by worker-worker transfer of a primer pheromone that inhibits foraging activity [6, 7]. The specific tasks workers perform are also strongly influenced by feedback from the behaviour of other workers and the environment. Foraging rates are regulated by the time foragers spend searching for food-storer bees that unload and store nectar, where short search times result in the recruitment of more foragers [8]. Thus, the more successful a forager is in collecting and unloading food, the more likely she is to continue foraging. Nonetheless, foragers with high daily flight activity exhibit a reduced maximum flight duration, and vice versa [9]. In bumble bees (*Bombus* sp., Hymenoptera: Apidae), which are primitively eusocial with a pronounced size variation in workers, task allocation appears to be more complex. Bumblebee workers do not undergo the same age-dependent changes in JH levels that honey bee workers do [10]. This could explain why their in-hive activities appear not to be age-specific [11, 12], except for callows (< 3d old) that frequently incubate brood [13]. It has been shown that *B. impatiens* workers are more inclined to continue a task they were performing the previous day than any other task, which is especially pronounced in guarding [12]. Other studies describe size-dependent task allocation (alloethism) in *B. agrorum* and *B. terrestris* [14–16]. For example, small bumble bee workers frequently engage in nursing activities, whereas large workers frequently take on guarding and fanning tasks [12, 14]. Larger workers also tend to forage more frequently, whereas smaller workers tend to remain inside the nest. One reason for this could be that large foragers are more efficient in collecting nectar per unit time than small foragers [16]. Furthermore, the body size / foraging range hypothesis predicts that a larger body size is advantageous as this enables bees to forage over longer distances [17, 18]. For example, there is evidence for a positive association between body size and maximum flight distance among stingless bee species [18] and within the species *Melipona mandacaia* (Hymenoptera: Meliponini) [17].

Although bumble bees have a relatively large body size, they are traditionally described as clumsy flyers due to their comparatively small wings. Nonetheless, they can forage under difficult ambient conditions such as cooler temperatures [16, 19], high altitudes [20, 21] and poor light conditions [22]. Astonishingly, *B. impetuosus* males can perform hovering flights under extremely reduced barometric pressures, mimicking conditions similar to altitudes surpassing the Mt. Everest [20]. Foraging flights of *B. terrestris* have been tracked with a harmonic radar up to 631 m away from their nest, but this method has a maximum tracking range of 700 m [23]. Microsatellite studies estimated foraging distances of more than 500 m on average to a maximum of almost 3 km in *B. terrestris* [24, 25]. An even higher potential average and maximum foraging range of about 2-5 km and 12 km respectively has been estimated for *B. vosnesenskii* in late season blooming clover fields [26]. Besides variation of foraging distances among species, there is considerable variation within species due to environmental conditions such as the location of the nesting sites [24, 27]. In *B. terrestris*, flight performance has been shown to be influenced by ambient temperature and worker body size, where 25 °C is optimal and larger workers can fly longer distances [19]. Thus, flight and foraging distances vary considerably between species, body size and environmental conditions.

Foraging comes with a considerable cost, as it can be risky, and more importantly, flying is a highly energy-demanding activity. Metabolic rates (MR) are about ten times higher during flight compared to terrestrial locomotion (Dickinson 1995, Harrison 2000) and 50-200-fold higher than MR at rest (Kammer 1978, Dudley, 2000). Honey bee foragers exhibit extremely high MR ranging between 100 to 120 mL O_2_ g^-1^ h^-1^, which is three time higher than the MR in hovering hummingbirds [28] and 50% higher than that of in-hive bees [29]. Elevated flight metabolism also leads to increased levels of reactive oxygen species (ROS), which are thought to have adverse effects on workers lifespan [30]. In fact, foraging activity has been found to be negatively associated with longevity in *A. mellifera* [31–33] and *B. agrorum* [15]. Non-foraging older and larger *B. impatiens* workers which develop larger oocytes, tend to remain inside the nest during the ergonomic colony phase[34]. This could be a strategy to avoid risky or energy-intensive tasks and likely preserves fat stores to increase their own reproductive success [34].

We hypothesised that (1) flight performance (distance and speed) of workers should increase with increasing body size, (2) flight performance should increase with age in smaller workers, (3) flight performance should decrease with age in large workers, and (4) older workers should have increasing relative fat reserves with increasing body size. As it is challenging to study flight performance in field or semi-field conditions, we measured flight performance in *B. terrestris* workers at three ages (i.e. 7, 14 and 21 days) under laboratory-controlled conditions using tethered air flight mills [19, 35, 36].

## 2. Materials and Methods

### 2.1 Experimental overview

The experiment was designed based a pilot study, in which the flight performance of differently aged workers from two colonies had been examined in the previous year. In this study, we repeatedly measured the individual flight performance of 58 workers from four colonies (n = 15, 18, 8 and 19 workers per colony) at the age of 7, 14 and 21 days. We selected workers with a wet body mass ranging from 100 mg to 320 mg, primarily during the ergonomic growth phase of each colony.

### 2.2 Bumble bee husbandry

We used four commercial, queenright *Bombus terrestris* colonies (Natupol Research Hives, Koppert B.V., Netherlands) with approximately 20-30 workers. They were housed in their standard plastic nest boxes (27 [l] x 24 [w] x 14 [h] cm). Each colony was connected to a small foraging arena (60 x 40 x 28 cm), where bees had *ad libitum* access to 40% w/v sucrose solution from 50 mL gravity feeders. Additionally, colonies were provided with 3-6 g of pollen candy (2:1 honey bee collected organic pollen: 75% sucrose syrup) daily, depending on their consumption. Colonies were kept under laboratory conditions with a relative humidity of about 40 % RH and an average room temperature of 23 ± 1 °C. Foraging areas were illuminated by two flicker free daylight-like LEDs (each 2400 lm, CRI98, 5500 K, True-Light International GmbH, Germany) under 14:10 h light:dark regime, but nest boxes were kept dark by covering them with cardboard.

### 2.3 Tagging and marking

Newly emerged workers (< 1 d old) were collected from their colonies. Their sex was determined by counting antennal segments (females have 12, males 13), under a stereo microscope. Then bees were immobilized with metal pins without harming them to carefully shave off the thorax hairs between the tegula, where a circular stainless-steel tag (0 2 mm, thickness = 0.1 mm) was attached using superglue (Supergel, UHU GmbH & Co. KG, Germany). These metal tags were colour coded to differentiate cohorts. To identify each individual bee, the middle legs were colour coded using water-based paint markers (5M Uni-Posca, Mitsubishi pencil, Japan). Afterwards, tagged bees were placed in a separate plastic cup for 30-60 min before returning them to their natal colony.

Although precautions were taken to guarantee that attached tags would not interfere with wing movements, tag positions were measured using a digital microscope (figure S1*a*, CHX-500F, Keyence GmbH, Germany) on frozen specimens at the end of the experiment. The tag deviation from the centre between the tegulae was calculated, but tag position did not significantly affect their flight performance (figure S1*b,c*).

### 2.4 Flight mill setup

Four tethered air flight mills like setups previously described were used (figure S2) f. The core of each flight mill is a lightweight and counter-weighted arm (length 32 cm) that floats by magnetic levitation and a needle that is inserted into a low-friction Teflon bearings at the centre of the arm. Individual bees were attached to a magnet (Ø 2 mm, 4 mm long) on one end of the arm and counter-balanced on the other arm, enabling tethered flights with their own power. A Hall effect magnetic sensor transmitted a voltage pulse every half rotation (flight distance of 50 cm), recorded to a PC using the software *guiBee* [37]. Data extraction and calculations of flight distance, duration, and speeds were executed using the RScript *FlightMillDataExtraction* [38] in R (version 4.3.0). Each flight mill was positioned at the centre of a plastic cylinder (Ø 46 cm), keeping about 7 cm distance between the bee and the cylinder wall. The inside was decorated with 2.5 cm wide black and white vertical stripes continuously printed on paper to provide consistent visual feedback [39]. The walls also prevented interference from neighbouring flight mills and reduced any potential impacts of air currents[40]. The flight mills were illuminated by four flicker free daylight-like LEDs (each 2400 lm, CRI98, 5500 K, True-Light International GmbH, Germany) from a height of 70 cm.

### 2.5 Flight trials

Marked workers aged 7, 14 or 21 days were gently collected from each colony in the morning using tweezers. They were kept separately per colony in a metal cage (9.5 x 8.5 x 5 cm) with *ad libitum* access to 40% w/v sucrose solution. After collection, workers were individually separated into flat-bottom glass vials (10 mL, 50 x 22 mm) with mesh lids, containing a 45 x 15 mm piece of cardboard to absorb any faeces. For 20 min each bee was individually fed with 40% w/v sucrose solution to satiation through the mesh of the lid. Subsequently, individual bees were weighed (d = 0.1 mg, analytic balance A210P-OD1, Sartorius GmbH, Germany) and attached to the magnet on the one side of the flight mill arm and kept in place on a launch platform. A counterweight (to ± 10 mg) was attached on the other side of the flight mill arm. The bees were then allowed to calm down and rest in the dark for 20 min by covering each flight mill with thick cardboard. Flight was initiated by positioning the bee in the direction of flight and quickly removing the launch platform. When a bee stopped flying, it was allowed to rest on a handheld plastic Petri dish (Ø 46 cm) for approximately 20 s. Each bee was allowed 4 stops, i.e. 5 flight starts. The temperature during all flight was recorded at 5 min intervals (RC-5 temperature data logger, Elitech Ldt., UK) to calculate the average flight temperature and account for slight room temperature differences. At the end of each flight trial, bees returned to their natal colonies. After the last flight trial at age of 21 d, bees were frozen and stored at −20°C until further analysis.

### 2.6 Measuring intertegular distance, dry mass, and fat content

In addition to the evaluation of the tag position, the intertegular distance (ITD) for each bee was measured using a digital microscope (figure S1*a*, CHX-500F, Keyence GmbH, Germany). This serves as an additional proxy for workers body size (figure 3a) [41]. Prior to measuring their dry weight and fat content [42], the sternites (ventral abdominal segments) of each individual bee was cut open from the stinger to the fourth sternite without damaging their guts. Then, bees were individually dried at 60°C for 3 d (drying cabinet U40, Memmert GmbH & Co. KG, Germany) and then weighed (d = 0.1 mg, analytic balance M-Pact AX224, Sartorius GmbH, Germany). To extract the body fat, bees were placed in 5 mL petroleum ether for 5 d. After discarding the ether and rinsing them with fresh ether, bees were dried for another 3 d and subsequently weighed. The fat contented was calculated as the difference between the dry weight and the dry weight after fat extraction.

### 2.7 Statistical analyses

All statistical analyses and data visualizations were performed using R version 4.3.2 [43]. To analyse the effects of the fixed factors age and body size (using ITD as a proxy, see figure 3a) as fixed factors on the response variables flight distance, average and maximum flight speed, generalized linear mixed effect models (GLMMs) were ran using the *glmmTMB* package. Flight distances were log-transformed using log_10_ (x + 1) to improve model fit based on gaussian data distribution. To account for repeated measures of individual bee flight performance, Bee ID was included as a random factor. The covariates colony (figure S4) and temperature during flight (figure S3) were additionally included as random factors. Significance (p < 0.05) of model terms was determined using the *Anova* function of the *car* package [44]. Pairwise comparisons between age groups were conducted using the function *emmeans* [45] with Bonferroni correction. The individual flight improvement was analysed using *chisq-test* function to perform a Pearson’s *χ*^2^ test for count data with Yate’s continuity correction. The *cor.test* function was used to further describe the relationship between the ITD and flown distance at each age class. Model selection was performed based on the Akaike information criterion (AIC) and likelihood ratio tests. The final models were compared with their respective null-models. Model assumptions and dispersion of the data were checked using the *DHARMa* package [46]. Linear mixed effect models (LMMs) were used to assess the relationship between individual mass-specific fat content and flight performance at the age of 21 days, using *lmer* function of the *lme4* package [47]. The full model included ITD and relative body fat contend as fixed factors and flight temperature as random factor. Model assumptions were checked visually and met expectations.

## 3. Results

Flight distance was primarily influenced by worker’s age (*χ*^2^ = 65.90, df = 2, p < 0.0001; figure 1a,b). Pairwise comparisons revealed significant differences between 7-day-old and 14-day-old bees (Tukey: t-ratio = −7.95, p < 0.0001), 7-day-old and 21-day-old bees (Tukey: t-ratio = −5.33, p < 0.0001), and 14-day-old and 21-day-old bees (Tukey: t-ratio = −2.77, p = 0.019). Although 86% of the same bees improved in their flight capacity from the age of 7 to the age of 14 days, 62 % of workers worsened again at the age of 21 days (*χ*^2^ = 26.69, df = 1, p < 0.0001; figure 1b). Although the worker’s body size also influenced flight distance (*χ*^2^ =9.75, df = 1, p < 0.001; figure 1c), the effect was not as pronounced as that of age. Although there was no relationship between ITD and log-transformed flight distance detected at the age of 7 days (Person’s correlation: r = 0.1568, p > 0.05), there were weak correlations at the age of 14 days (r = 0.26, p = 0.05) and 21 days (r = 0.38, p < 0.01) respectively (figure 1c).

**Figure 1.**
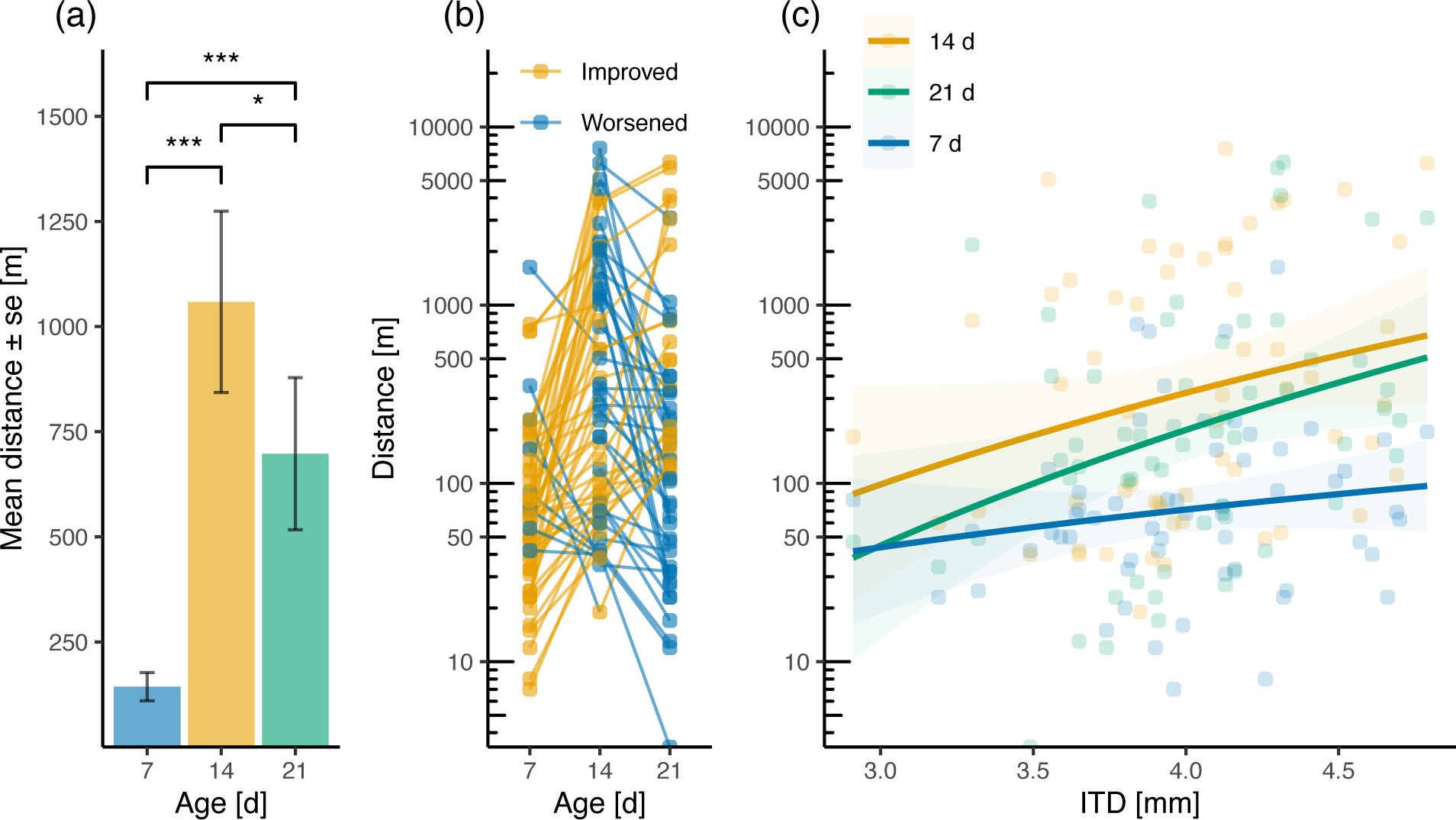
Effects on flight distance. (a) Effect of worker’s age in days on mean flight distance ± se [m]; (b) individual improvement in flight distance [m] as workers aged (yellow = improved, blue = worsened); (c) effects of ITD (proxy for body size) on flight distance [m] as workers aged (age classes: blue = 7 d, yellow = 14 d, green = 21 d old). In panel (a) and (b), the asterisks (*) indicate significant differences revealed from the post hoc test, with * p < 0.05 and *** p < 0.001. In panels (b) and (c), respectively, distance flown have been plotted on log10 scales as distances have been log-transformed for statistical analysis.

Average flight speed was affected by worker’s body size (*χ*^2^ = 17.99, df = 1, p < 0.0001; figure 2 b), but not their age (*χ*^2^ = 4.70, df = 2, p > 0.05; figure 2a). Similarly, maximum flight speed was also rather a result of worker’s body size (*χ*^2^ = 22.48, df = 1, p < 0.0001; figure 2b) than their age (*χ*^2^ = 6.84, df = 2, p < 0.05; figure 2*a,b*). Pairwise comparisons of slopes with maximum speed as response variable revealed a steeper and higher flight distances in 14-day-old bees compared to 7-day-old bees (Tukey: t-ratio = −2.61, p < 0.01), but no differences between 7-day-old and 21-day-old bees (Tukey: t-ratio = −1.19, p > 0.05) nor 14-day-old and 21-day-old bees (Tukey: t-ratio = 1.44, p > 0.05). Nonetheless, ITD correlated only weakly with mean speed at the age of 14 days (r = 0.33, p < 0.05) and 21 days (r = 0.49, p < 0.0001), but there was again no correlation for 7-day-old workers (r = 0.16, p > 0.05) respectively (figure 2*b*).

**Figure 2.**
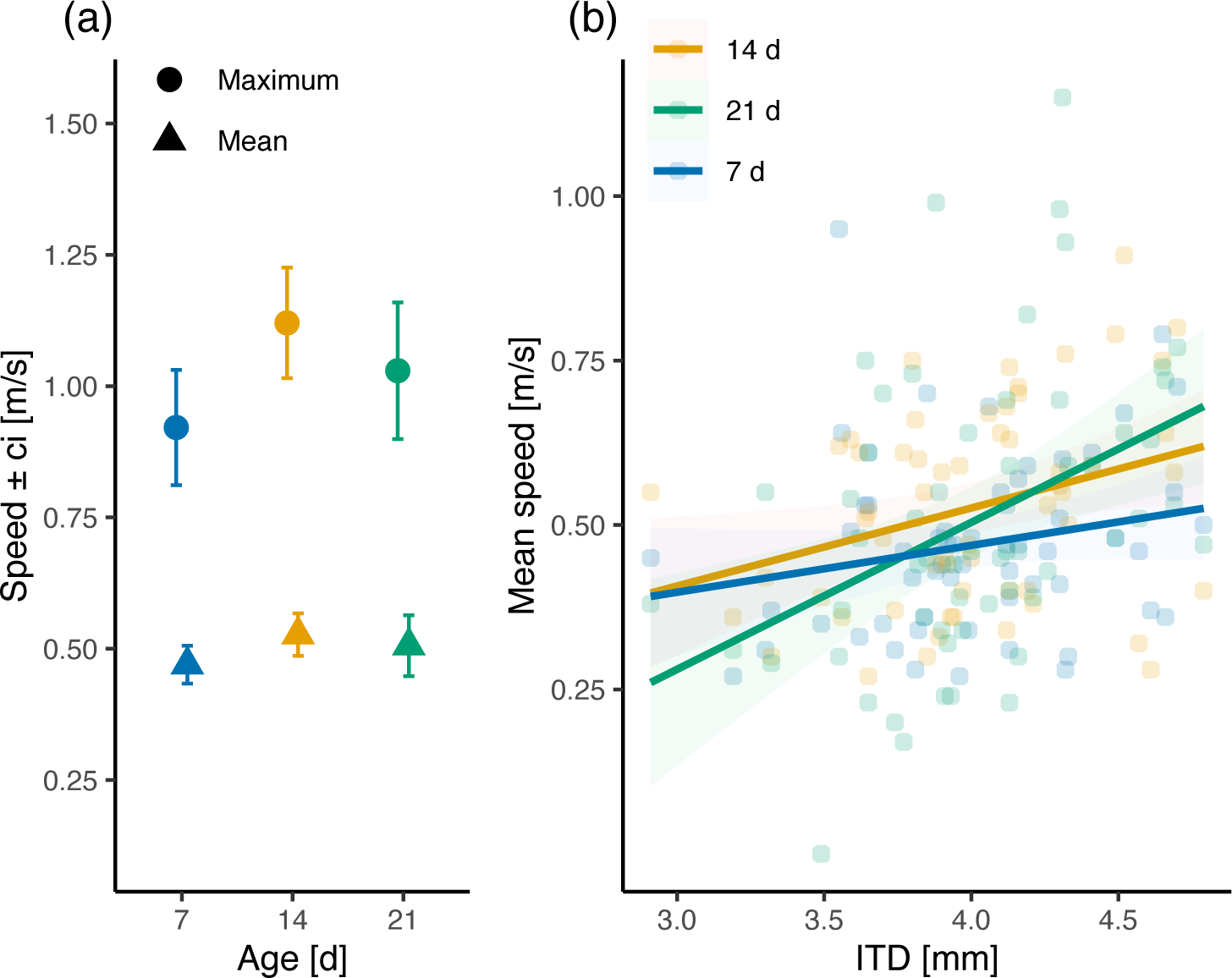
Effects on flight speed. (a) Effect of worker’s age in days on mean (triangle) and maximum (circle) flight speed ± ci [m/s]; (b) effects of ITD (proxy for body size) on average flight speed [m/s] as workers aged (age classes: blue = 7 d, yellow = 14 d, green = 21 d old). Asterisks (*) indicate significant differences revealed from the post hoc tests, with * p < 0.05, ** p < 0.01 and N.S. indicating p > 0.05.

The analysis of the morphometric data of 21-day-old workers revealed that mass-specific fat content decreases with increasing ITD (Person’s correlation: r = - 0.70, p < 0.0001, figure 1*a*, green circles), while dry mass was strongly positively correlated with ITD (r = 0.78, p < 0.0001, figure 1*a*, grey circles). There was a significant effect of ITD, mass-specific fat content, and their interaction on the log-transformed flight distance in 21-day-old workers (LMM: *χ*^2^ = 17.28, df = 3, p < 0.001). This effect was explained by ITD (*χ*^2^ = 4.93, df = 1, p < 0.05), while neither relative fat content (*χ*^2^ = 0.09, df = 1, p > 0.05, figure 3*b*) nor their interactions (LMM: *χ*^2^ = 0.37, df = 1, p > 0.05) showed a significant influence. Similarly, ITD and mass-specific fat content and their interaction significantly affected flight speeds in 21-day-old workers (LMM: *χ*^2^ = 13.15, df = 3, p < 0.01), which was explained by ITD (*χ*^2^ = 8.98, df = 1, p < 0.01) and not relative fat content (*χ*^2^ = 1.11, df = 1, p > 0.05, figure 3*c*) and their interactions (*χ*^2^ = 0.36, df = 1, p > 0.05).

**Figure 3.**
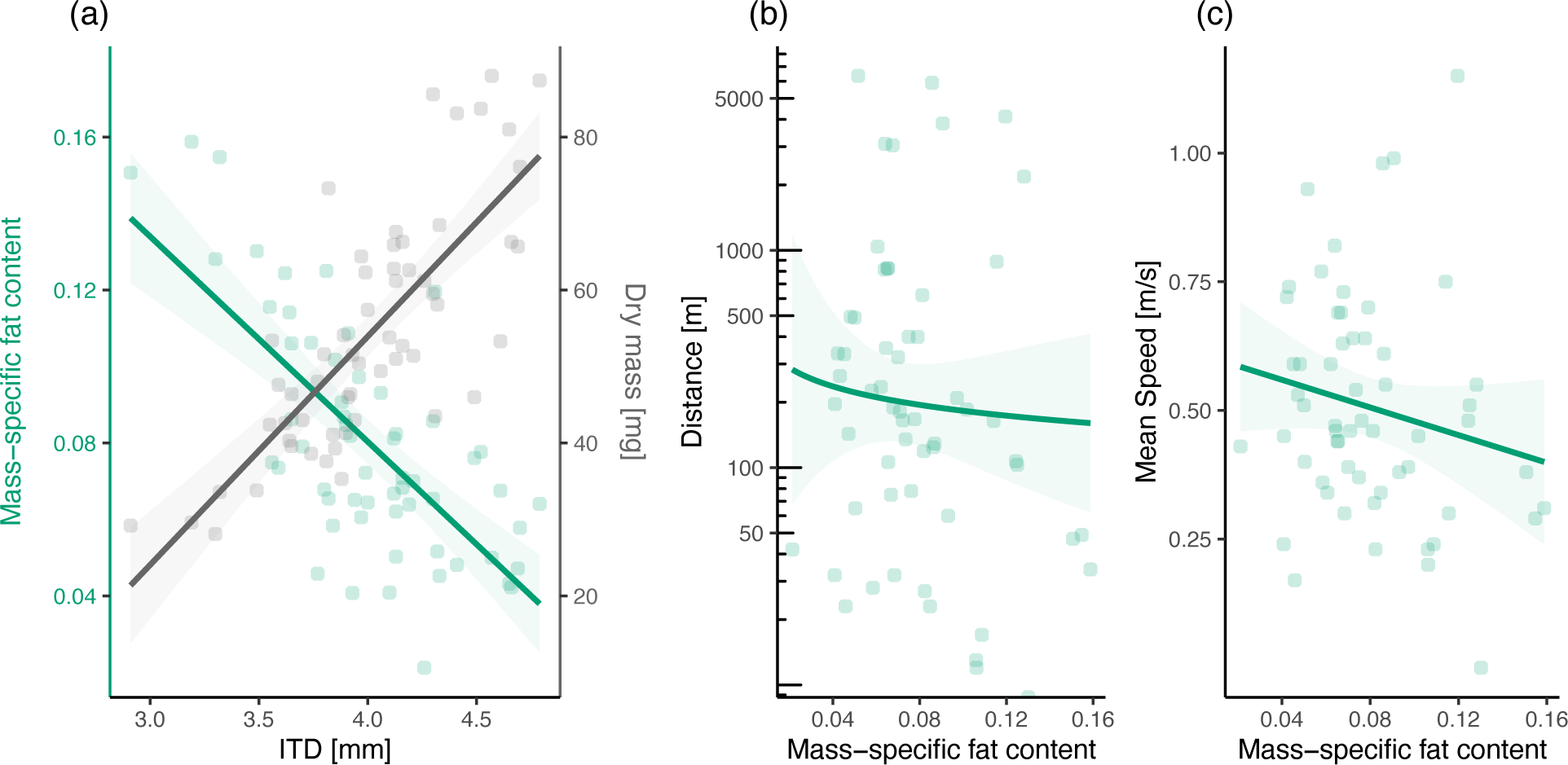
Associations of mass-specific fat content of 21-day-old workers. (a) Intertegular distance (ITD) positively correlated with dry mass (grey circles, Person’s correlation: r = 0.78, p < 0.0001) and negatively with relative fat content respectively (green circles, r = - 0.70, p < 0.0001), respectively. There was no correlation between relative fat content and log_10_ flight distance (b, r = - 0.20, p > 0.05) and mean speed (c, r = - 0.19, p > 0.05).

## 4. Discussion

Our data provide clear evidence that older bees fly longer distances in *B. terrestris* workers. Several studies have explored the variance in task allocation, foraging behaviour, foraging efficiency, or flight ability among bumble bee workers, attributing these to their pronounced size polymorphism [12, 14–16, 19, 22, 48–55]. This explanation is also consistent with the hypothesis that larger foragers can fly longer distances as described in stingless bees[17, 18]. For example, large foragers of *M. mandacia* have an up to 24% larger flight range and 48% higher maximum foraging distance compared to small foragers[17]. Some studies also describe that the age at which bumble bee workers change to out-hive tasks varies substantially and some workers may never forage [11–15].

In our experiment, most bees (70%) flew less than 100 m at the age of 7 days (figure 1). This is significant as previous studies on, for instance, the influence of ambient temperature or pesticide exposure on flight performance, used 100 m flight distance as a threshold for flight motivation [19, 56]. Whether bees flew more than this was independent of body size (figure 1*c*). This contrasts with findings indicating that the transition to foraging occurs in large bumble bee workers at the age of 5 days, while small workers delay this transition until the age of 15 days[15]. Flight performance, or motivation, improved in 85% from the first test (7 days) to the second (14 days) (figure 1*b*). In fact, they flew on average more than six times further (figure 1a) and with a 22% increase in maximum flight speed (figure 2*a*). Despite average and maximum flight speeds being influenced by age (figure 2*a*), our data suggest that flight speeds are rather body size-dependent (figure 2*b*). Hence, our results support previous data indicating that larger forager tend to fly faster [50].

The transition to foraging has been associated with elevated expression levels of the so-called *foraging* gene (*for*) that encodes the cGMP-dependent protein kinase (PKG) [57]. Honey bee foragers exhibit higher *Amfor* gene expression along with increased PKG activity compared to nurses [58]. Similarly, it has been shown that *Btfor* is expressed more in larger than in smaller *B. terrestris* workers [53]. This suggests that larger workers rather tend to forage as PKG plays an important role in muscle cell proliferation [57]. Flight muscles typically constitute 12% to 16% of an insect’s body mass, and an increase in flight muscles can result in improved flight ability [59]. About 50% of the variability in the MR in flight muscles of bumble bees is explained by body size and about 75% when accounting for wing morphology and enzyme activity [60]. This would support our observed size-dependent flight speed (figure 2*b*). While honey bee workers lose 42% of their wet body mass in the transition to foragers [29], we did not find such age-dependent weight loss (figure S3*b*). We speculate that our age and size-dependent pattern indicate that bumble bees must first built up or train their flight muscles before being capable of flying longer distances as has been shown in male *Plathemis Lydia* dragonflies (Odonata: Libellulidae) [61].

Our data further suggest that flight performance peaks around the age of 14 days, after which 62% of workers worsened in performance again at the age of 21 days (figure 1*b*). A decrease in *Btfor* expression levels has also been described in aging bumble bee foragers[53]. This would be consistent with the age-dependent metabolic properties and flight performance observed in honey bees. It has been shown that foraging activity and flight capacity peaks between the age of 15 and 32 days in honey bee workers depending on the season [29, 62]. As insect flight is metabolically extremely costly, another explanation for this pattern could be the necessity to acquire and energy reserves. For instance, honey bee foragers exhibit the highest glycogen reserves in their flight muscles, acting as a buffer against starvation and aging, which diminish as they age [9]. Nonetheless, the primary energy reservoir to power bee flight is trehalose in the haemolymph [63–65].

The mean trehalose concentration is more than twice as high in foragers than in nurses and newly emerged honey bee workers [66]. Although the energy demand in bumble bees remains constant and independent of flight speeds ranging from hovering to 4.5 ms^-1^ [67], it is possible that trehalose reserves vary considerably in bumble bee workers. Although little is known about the haemolymph sugar concentrations in bumble bees, such variability has been reported among returning foragers, potentially depending on environmental factors such as flower availability[68]. For *B. terrestris,* the foraging distance from the nest ranges from an average of 500 m to a maximum of 3 km [24, 25], which aligns well with our measured flight distances in 14-day-old and 21-day-old workers (figure 1). It has been observed that bumble bee workers forage at some distance to their colony [23, 69]. One potential reason for this could be the avoidance of competition with other bumble bees or honey bee colonies [68].

As high flight activity leads to reduced longevity [15, 30–33], this could also affect the reproductive output of worker bumble bees. While it has been suggested that larger workers avoid energy-intensive tasks as they age to reserve fat stores for oocyte development [13, 34], we did not find evidence that larger workers fly shorter distance or with lower speeds as they age (figures 1*c* and 2*b*). Moreover, our data indicate that with increasing body size, mass-specific fat stores decrease in 21-day-old workers (figures 3*a*). Regardless, mass-specific fat stores did not significantly influence flight distance nor flight speed at the age of 21 days (figure 3*b,c*). It could be that the colony phase plays an important role in this process [11, 13, 33, 34]. For example, it has been shown that workers of *B. bifarius* begin foraging earlier as the colony matures, which could explain decreased longevity in bumble bee workers at a later colony stage [11].

In conclusion, our data indicate that it is primarily age, not body size, that influences bumble bee workers’ flight distance. This finding contrasts the body size foraging range hypothesis. Their pronounced body size variation, however, influenced flight speeds in our experiment as well – just to a lesser degree than age. As bumble bees have become an important model system for wild bees for studying, for example, their response to climate change, the effects of pesticides, and evolutionary ecology and cognition more broadly, we suggest that accounting for worker’s age besides body size should be carried out as standard when conducting experiments.

## Data accessibility

All data and statistical code output supporting the findings of this study are available from the Dryad Data Respository [Anonymous, 2024].

## Author’s contributions

MG carried out the experimental work and gave approval for publication. CK conceived and designed the study, assisted with the experimental work, analysed and interpreted the data, and drafted the manuscript.

## Acknowledgments

We thank Carolin Bäuml and Helena Schulte for their assistance in the laboratory. Our special thanks to Günter Rödl and Johann Schmid from our institute’s mechanical and electronic workshops for their invaluable help in constructing the flight mills. Lastly, we thank Dr. Tomer J. Czaczkes and Prof. Dr. Erhard Strohm for their helpful comments on an earlier version of this manuscript.

## Supplementary figures

**Figure S1.**
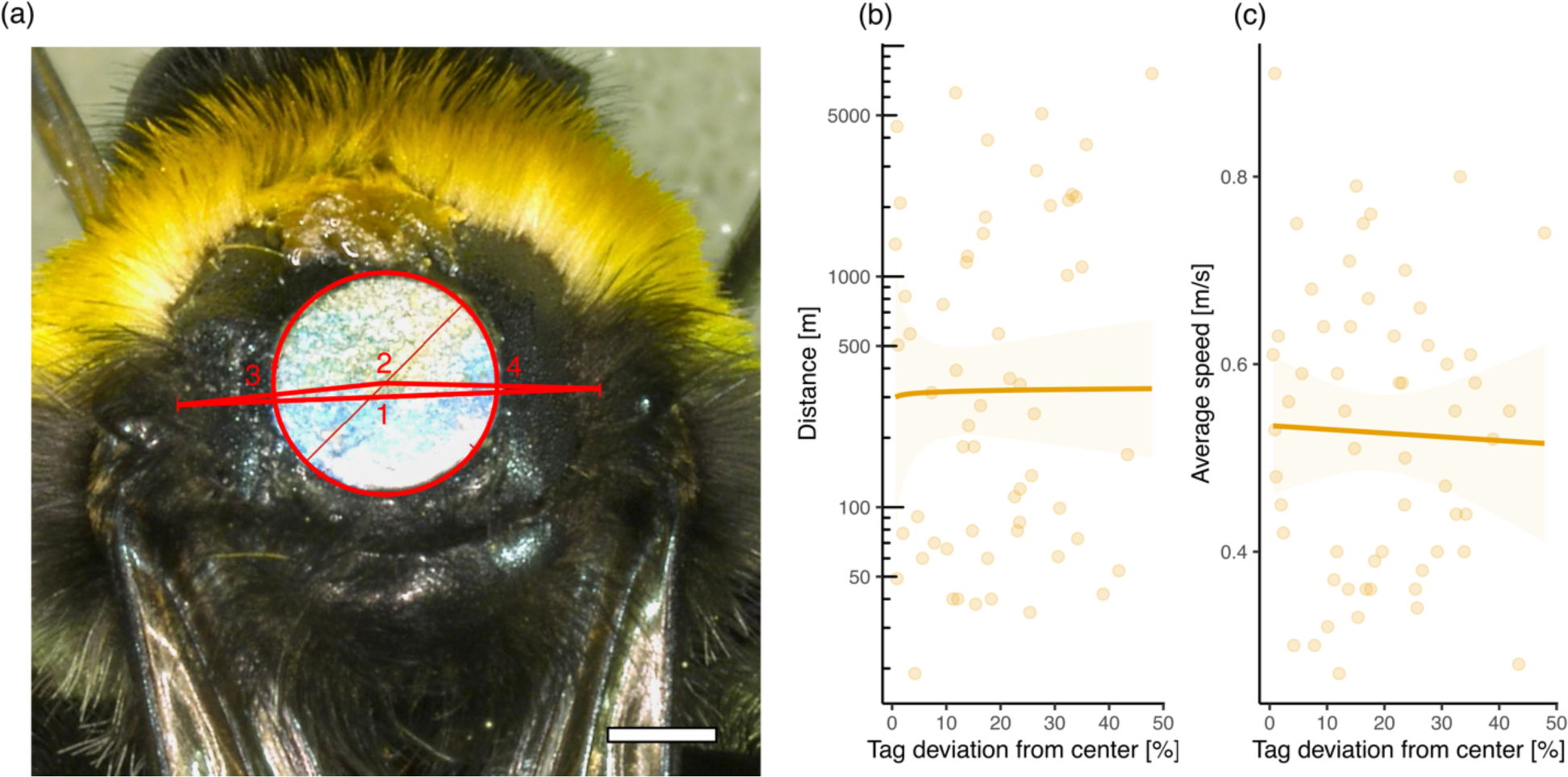
Measuring intertegular distance (ITD) and evaluation of tag position. (a) Relative tag deviation was determined by calculating the absolute difference between distances from the tegulae (3 and 4) to the centre point of the metal tag (2), divided by distance between tegulae (1). Scale = 1 mm. There was no significant influence of the relative tag deviation on (b) flight distance and (c) average flight speed.

**Figure S2.**
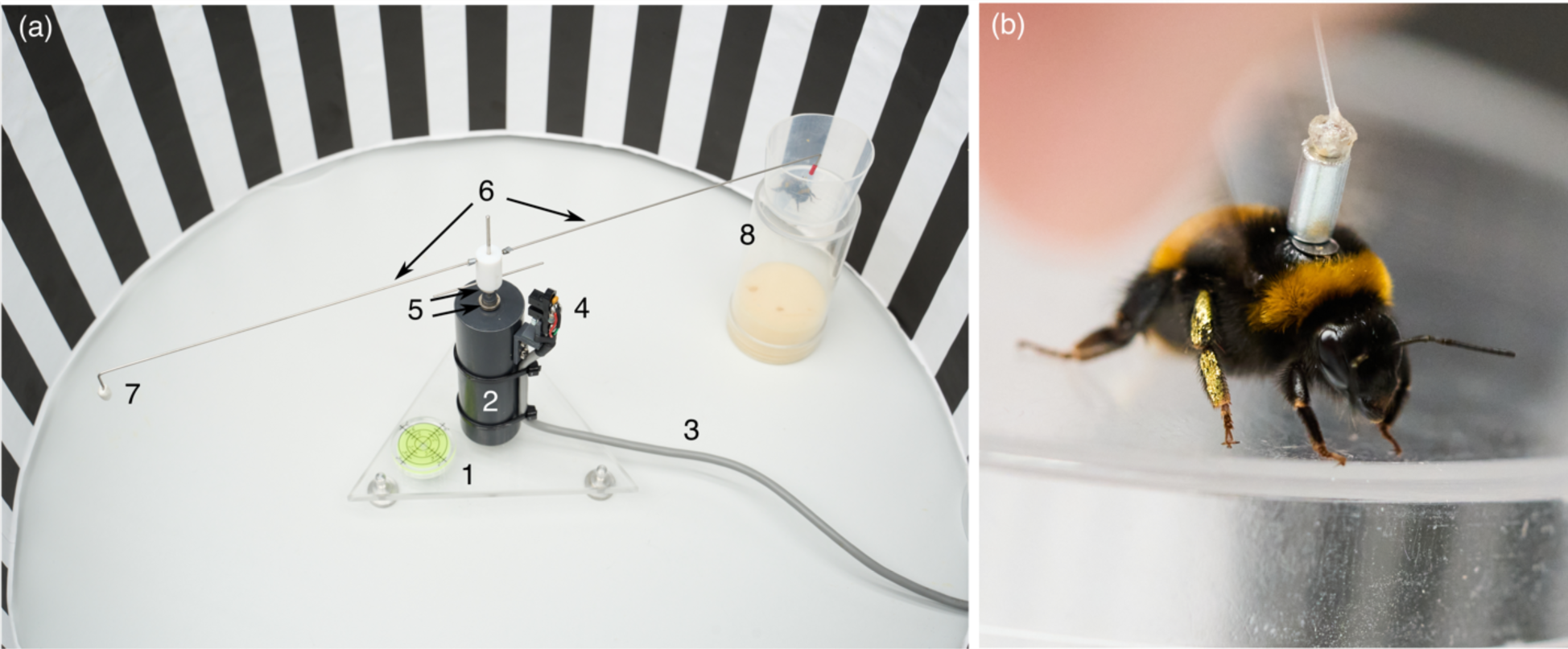
Tethered air flight mill setup. Each flight mill consisted of (a) a height-adjustable base with a bubble level (1); Delrin role (2); USB cable (3) connecting Hall effect magnetic sensor (4) to a PC; opposing magnet rings (5); flight mill arms (6), which were attached on a low-friction Teflon bearing; one arm held an adjustable counter weight (7) and the other arm held a small magnet, where tagged bees were attached (b). Prior to flight trials bee rested for 20 min in a supporting stand (8) in the dark. Flight mills were position at the centre of plastic cylinder (diameter = 46 cm) with black and white vertically striped (width = 2.5) walls.

**Figure S3.**
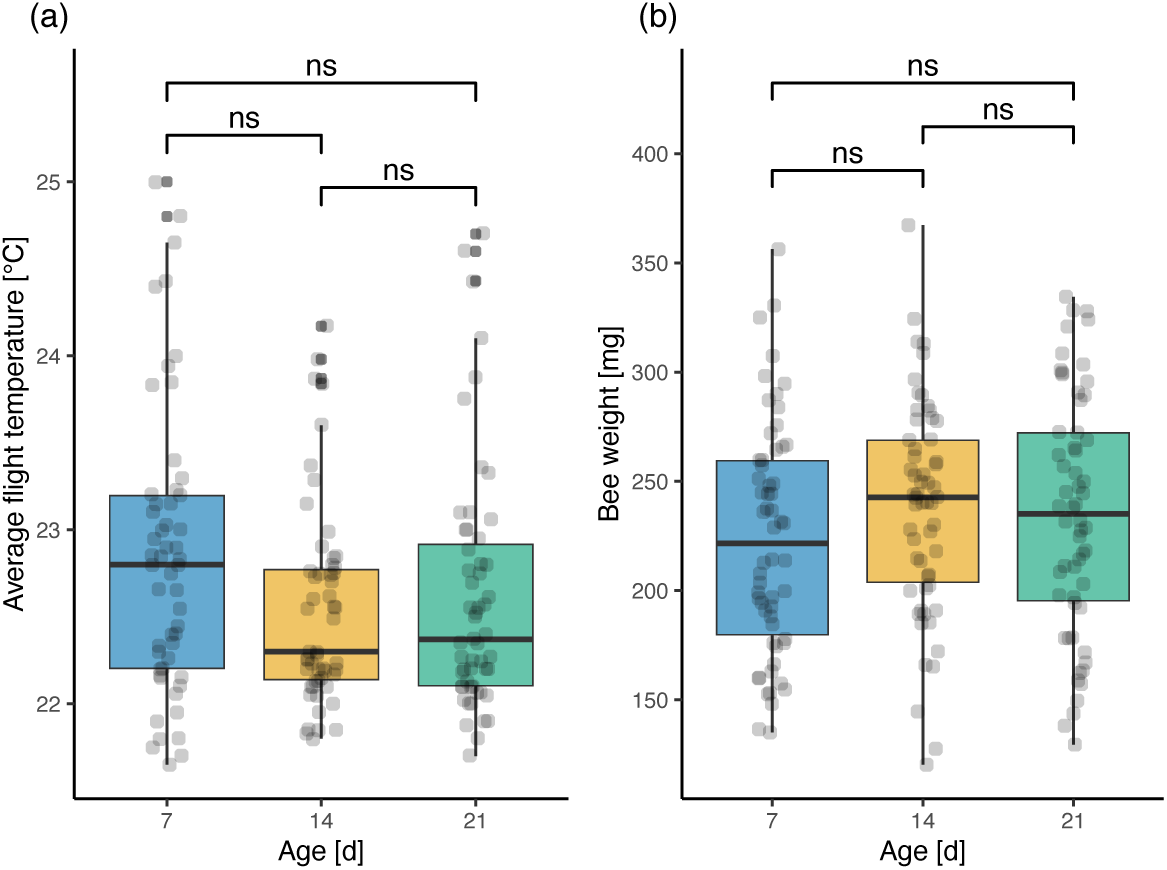
Comparisons between age classes for co-variates (a) average flight temperature and (b) bee wet weight before flight trials. There was no significant difference between age classes and (a) average flight temperature (Kruskal-Wallis rank sum test: *χ*^2^ = 5.21, df = 2, p > 0.05), and (b) bee wet weight before flight (ANOVA: F_2,171_ = 1.23, p > 0.05) respectively. Multiple comparisons of post-hoc tests (Wilcoxon ranks sum test and Tukey HSD respectively) between age classes were not significant (ns, p > 0.05).

**Figure S4.**
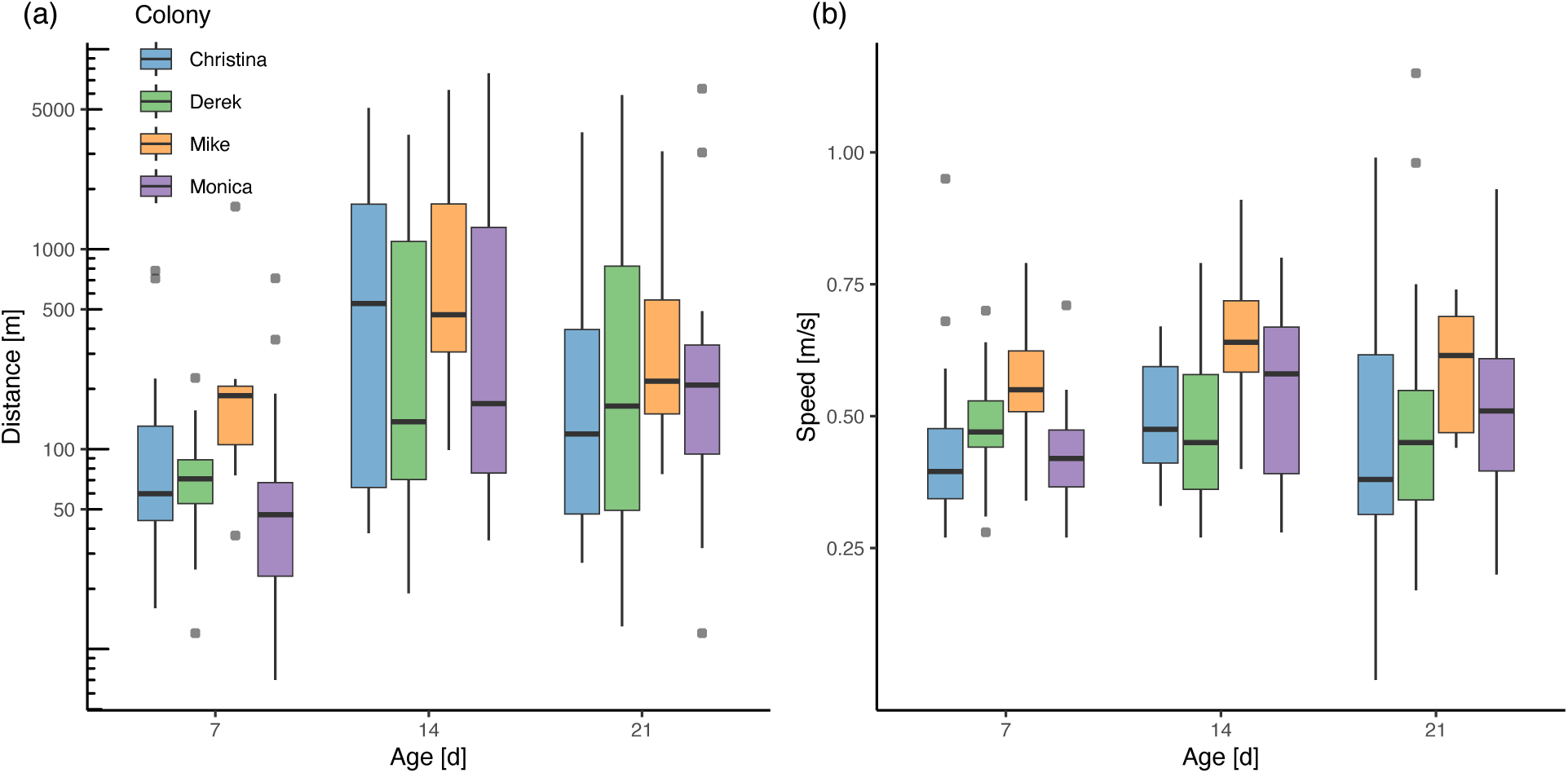
Data distribution for (a) flight distances and (b) mean flight speeds between per colony. (blue: Christina, n = 14; green: Derek, n = 17; orange: Mike, n = 8, purple: Monica, n = 19).

